# Isotopic evidence for population dynamics in the Central Italian Copper Age and Bronze Age

**DOI:** 10.1101/2021.09.30.462554

**Authors:** Marco Romboni, Ilenia Arienzo, Mauro Antonio Di Vito, Carmine Lubritto, Monica Piochi, Maria Rosa Di Cicco, Olga Rickards, Mario Federico Rolfo, Jan Sevink, Flavio De Angelis, Luca Alessandri

## Abstract

The mobility patterns in the Italian peninsula during prehistory are still relatively unknown. The excavation of the Copper Age and Bronze Age deposits in La Sassa cave (Sonnino, Italy) allowed to broaden the knowledge about some local and regional dynamics. We employed a multi-disciplinary approach, including stable (carbon and nitrogen – C and N, respectively) and radiogenic (strontium-Sr) isotopes analyses and the identification of the cultural traits in the material culture to identify mobility patterns that took place in the region. The Sr isotopic analyses on the human bones show that in the Copper Age and at the beginning of the Bronze Age, the cave was used as a burial place by different villages, perhaps spread in a radius of no more than 5 km. Stable isotopes analyses suggest the introduction of C4 plants in the diet of the Middle Bronze Age (MBA) communities in the area. Remarkably, in the same period, the material culture shows increasing influxes coming from the North. This evidence is consistent with the recent genomic findings tracing the arrival of people carrying a Steppe-related ancestry in Central Italy in MBA.

## Introduction

La Sassa cave (Sonnino, Latina province in Southern *Latium*) was explored since 2015 in multiple archaeological campaigns ^1,2^. The cave holds a multi-stratified archaeological archive from the Late Pleistocene to present days (Fig. 1). The cave area, subdivided in progressively numbered rooms, is articulated in at least 8 Rooms. The osteological material in La Sassa constitutes multiple bone assemblages reflecting naturally occurred arrangements of the pristine primary depositions, as several tiny bones originally tied in weak diarthroses, such as metacarpals and phalanges, were recovered. Human remains were found in three soundings (L, WD, and N) in Room 1 (Fig. 1, b). No diagnostic potsherds associated with these remains have been found, but the stratigraphy and the radiocarbon dates from human bones constraint these layers to the Copper Age (CA) and Early Bronze Age (EBA) ^3^. Some human remains have also been collected in Room 2. Again, no associated diagnostic potsherds were found, but radiocarbon dates fall in the CA ^3,4^. Few human bones in Room RA belong to an infant whose femur was radiocarbon dated to the Middle Bronze Age (MBA), sub-phase 2 ^3^.

**Figure 1:**
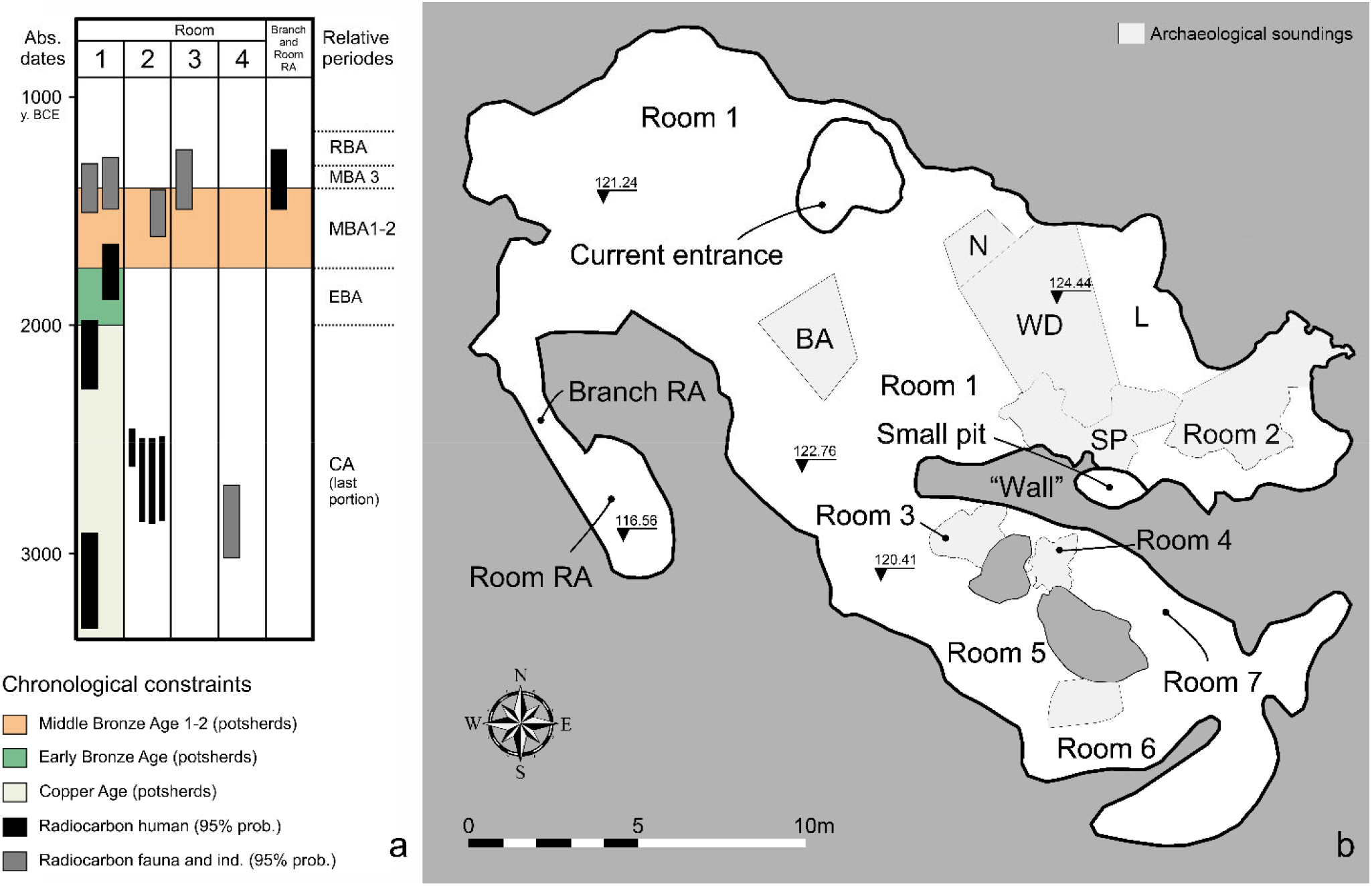
Chronology and soundings at La Sassa cave. a) Chronological constraints of the different rooms of the cave. CA, Copper Age; EBA, Early Bronze Age; MBA, Middle Bronze Age; RBA, Recent Bronze Age. No reliable radiocarbon dates are available for the boundary between MBA sub-phases 1 and 2 in Central Tyrrhenian Italy. The MBA subphase 1 can be further subdivided in 1A (earlier) and 1B (later) based only on pottery stylistic traits. b) Map of the soundings at La Sassa cave. Outline of the cave from photogrammetric survey (Rooms 1-2 and partly 3)^2, 5^, and instrumental survey (Rooms 4-7 and partly 3).

An extensive set of isotopic and preservation analyses has been performed on bones and teeth enamel to characterize individual mobility and dietary patterns of people buried in La Sassa. Soil and water samples, as well as faunal remains, were analyzed to obtain the local isotopic baselines. The results were put in the context of recent genomic and isotopic findings, suggesting the area surrounding the Pontine plain (also known as Agro Pontino) as a complex stage for the population dynamics in prehistoric Central Italy.

Most of archaeological items testifying for settlements of the CA around La Sassa were found by chance (Fig. 2). Some isolated items come also from the Lepini and Ausoni Mounts ^6^. Burial places have been recognized in two nearby caves: the Scalelle and Vittorio Vecchi caves. The burial goods collected at the Scalelle cave ^7^ parallelize with the southern Italian CA culture named *Gaudo* ^8^. The oldest radiocarbon dates for this southern culture in *Latium* ranges from 3330 to 3210 BCE ^9^.

**Figure 2:**
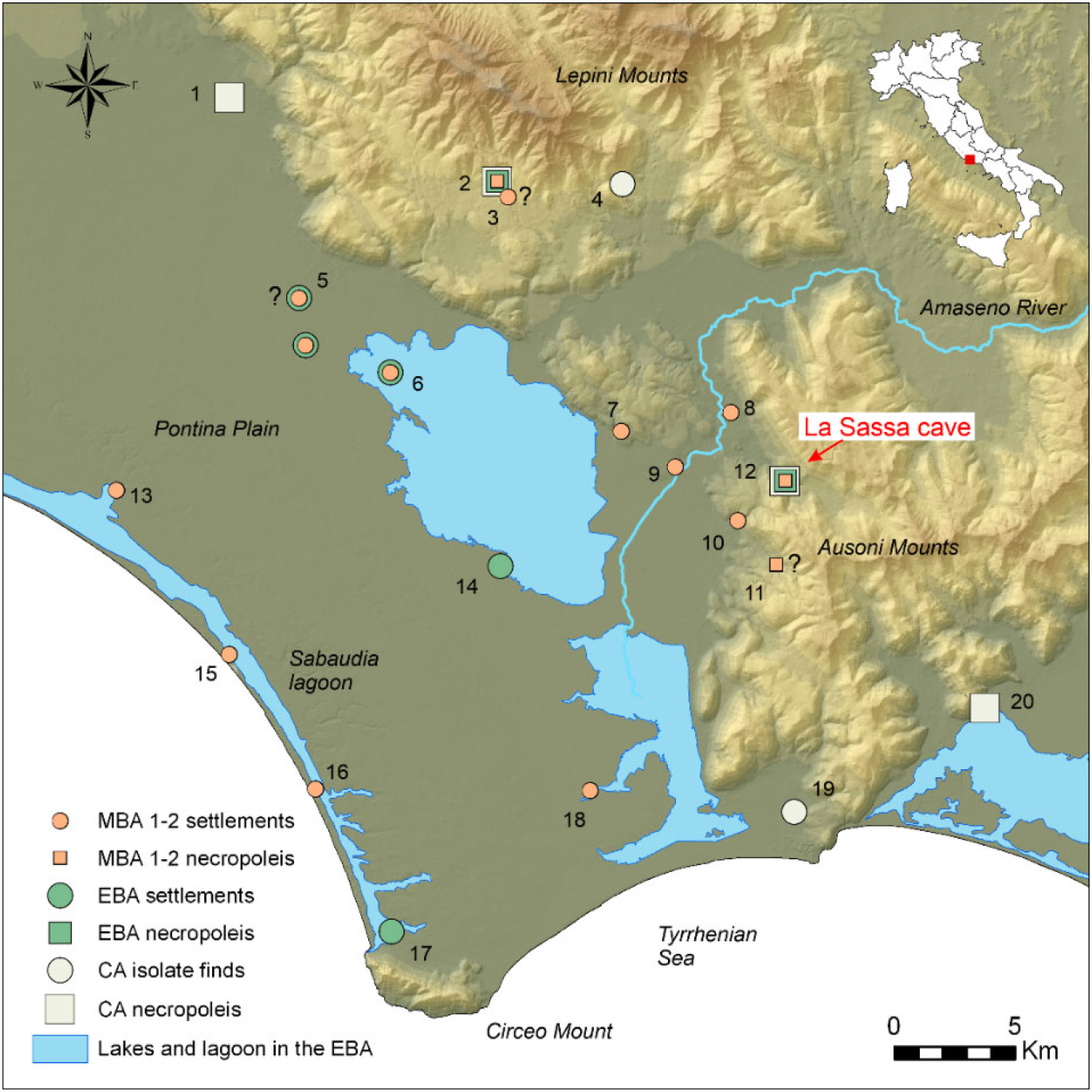
La Sassa cave on the Copper Age (CA), Early (EBA) and Middle Bronze Age (MBA) sub-phases 1-2 landscape. Entrance coordinates: UTM WGS84: 352628E, 4587449N, municipality of Sonnino. 1, Vacchereccia; 2, Grotta Vittorio Vecchi; 3, Longara; 4, Monte Acuto; 5, Tratturo Caniò; 6, Proprietà Ricci; 7, Valle Fredda; 8, Mola dell’Abbadia; 9, Colle Pistasale; 10, Colle Colanero; 11, Pistocchino cave; 12, La Sassa cave; 13, Colle Parito; 14, Mesa; 15, Caprolace; 16, Caterattino; 17, La Casarina; 18, Borgo Ermada; 19, Terracina (municipality); 20, Scalelle. The EBA from Tratturo Caniò and the MBA sub-phases 1-2 from Longara are uncertain. In the Pistocchino cave no archaeological excavation has been performed so far: the interpretation as necropolis is uncertain. Sites from Alessandri 2013; Anastasia 2007; Carboni 2002; Tol et al. 2021. Recostruction of the lakes and lagoons from Alessandri 2013; van Gorp, Sevink, and Van Leusen 2020. Background DEM from TINITALY/01 ^20^

In Vittorio Vecchi cave, hundreds of disarticulated human bones have been found together with abundant EBA and MBA potsherds, and a few CA potsherds ^10^. Despite the lack of radiocarbon dates, it is possible the narrowing of the funerary use of the cave to the BA, which starts between 2100 and 2000 BCE in the central Italian chronological framework ^11^. No EBA evidence has been found so far in the Lepini and Ausoni Mounts.

In the nearby Pontine plain, only a few finds have been collected ^12^: an isolated bronze axe in the Sabaudia lagoon ^13^, and some potsherds below the tephra of the Avellino eruption ^14^, recently dated around 1900 BCE ^11,15,16^. The lack of CA and EBA occurrence in the mountains might be due to the sparse unsystematic research in the area. In contrast, the paucity of evidence in the nearby Pontine plain seems to reflect a low population density in the CA. However, multiple sites arose along the coastline and at the foot of the Ausoni Mounts in the next MBA sub-phases 1 and 2 ^13^.

### Geological setting

La Sassa cave is situated in the west of the Ausoni Mounts, which form part of the Lazio-Abruzzi Meso-Cenozoic carbonate platform of Central Italy. These mountains consist of thickly bedded, relatively homogenous limestones dating from the Cretaceous to Paleocene (Carta Geologica d’Italia, sheet 159, Frosinone). The northern extension of the Ausoni Mounts is bordered by a series of elongated tectonic basins that form the Amaseno river catchment. The Amaseno first runs through a narrow valley in between the Ausoni Mounts and aeolian sand-covered limestone hills of Priverno/San Martino and then enters the Agro Pontino graben. There the Holocene deposits prevail, with some outcrops of older, lagoonal deposits of Eemian age ^21^, and alluvial fans and debris slopes descending from the Ausoni Mounts (more details in Supplementary, chapter 1).

### Evidence of regional mobility in the Italian context

Multiple pottery items were found in the local contexts. From a ceramic typological perspective, during the CA and the EBA, the area around La Sassa cave seems to be strongly influenced by (or even part of) the southern cultures of *Gaudo* (CA) and *Palma Campania* (EBA). At the beginning of MBA 1-2, the potsherds so far collected in the Pontine plain show influences both from the northern *Grotta Nuova* ^22^ and the southern *Protoappeninico* cultures ^23^. Indeed, the boundary between the latter had already been hypothesized in the Pontine plain ^23,24^. The comparison between the Vittorio Vecchi and La Sassa funerary contexts is worth exploring due to their liminal chronological and topographical settings. The MBA sub-phase 1A (MBA1A) potsherds recovered in La Sassa show *Protoappenninico* stylistic traits. Conversely, the MBA sub-phase 1B (MBA1B) potsherds show the first traces of *Grotta Nuova* styles. A mirror situation can be traced in the Vittorio Vecchi cave, where the MBA1A potsherds point out at *Grotta Nuova* styles, while in the MBA1B the first traces of *Protoappenninico* are noticeable.

These stylistic differences suggest that the local boundary between the two cultural contexts might be placed around the Amaseno valley (Fig. 2), and their contacts increased starting from the MBA1B. Recent genomic findings ^4^ confirm the area surrounding the Pontine plain as a liminal zone not only for cultural circulation but rather for mobility of people with different genomic characteristics that could have moved throughout that area in the BA. Indeed, even considering a few CA individuals from La Sassa cave, the signature of BA movement from genomically Steppe-related people has been demonstrated through Italy, emerging in the EBA in Northern Italy and moving southward through time ^4^.

So far, the in-depth analysis of the individuals buried in La Sassa appears pivotal in broadening the knowledge about the population dynamics occurring in that area at the CA/BA boundary.

To date, the mobility in Italian CA and BA communities has only been partially explored through Sr and O isotopic analysis. Recently, De Angelis and colleagues ^25^ found evidence of a sedentary lifestyle rather than extensive mobility in Central Italian CA burials, like pointed out in some Southern Italian sites ^26^. Conversely, the mobility practices of the northern Italian communities across a timeframe ranging from the EBA to the Late Bronze Age (LBA) suggested that people spurted their mobility ^27^.

### Carbon and nitrogen stable isotopes background in the Copper Age and Bronze Age

The analysis of carbon and nitrogen stable isotopes of bone proteins was widely advocated to investigate further ancient bio-cultural characteristics such as diet ^28^. Often, a change in diet might be suggestive for people moving from elsewhere carrying novel cultural and biological features. This theoretical framework was already applied to explain the dietary differences in multiple Italian BA funerary contexts ^29,30^.^29,30^ Indeed, recent surveys related to CA and BA communities in Central Italy underlined that this chronological and cultural transition represented a critical step for the Italian populations, characterized by economic changes and technological improvements ^29,31^. A terrestrial diet based on C3 plants typical of the ‘Neolithic package’ (wheat and barley) with a moderate animal protein intake was demonstrated for CA-EBA Italian communities. At the same time, new crops (C4 plant, i.e., millet) were progressively introduced into the Italian peninsula in the MBA, though their consumption might have been limited to occasional period or particular sites ^29,31,32^.

The new crop species seem to be inlet into Europe from the Eurasian Steppe regions during the Neolithic and introduced into the Italian peninsula in BA ^33,34^. The increased adaptability and frequent harvesting periods made the C4 as suitable crops for challenging environments. So far, the spread of the C4 plant consumptions seems to be restricted to Northern and Central Italy in the MBA. The southernmost fringe for their exploitation was represented by people buried in Misa cave and, possibly, Felcetone in the Latium/Tuscany boundary ^29^. Still, limited consumption of C4 plants was detected through starch grains identification in Scoglietto cave ^35^, and recently for a few diachronic samples in Abruzzi ^36^. However, it is not fully understood if these plants were consumed or constituted animal fodder. Whatever being the role for the C4 plants in Northern and Central Italy, the Southern areas of the peninsula seem to be generally grounded on moderate proportions of animal protein and C3 plants ^32^.

## Results

### Strontium isotopic results and FT-IR analyses

We determined the Sr isotopic compositions of spring water and soil samples in a radius of c.a. 5km around the cave to characterize the “local baseline” (Table S3). Soil samples are featured by Sr isotope ratios varying from 0.70983 (sampled close to Priverno) and 0.70799 (sampled within the cave). Sr isotope ratio determined for the cave soil is the lowest among those analyzed and similar to that of the microfauna tooth enamel (*Microtus sp*.). Moreover, three soils are isotopically similar to a limestone from Mt. Lepini (0.7088) ^37^ and to Miocene limestone sampled on the Aurunci Mountains (0.7085) ^38^. The remaining soil samples are enriched in radiogenic Sr and fall within the range of ^87^Sr/^86^Sr characterizing the volcanic products form the Middle Latin Valley ^37^. The domestic fauna tooth enamel values show a wide range of ^87^Sr/^86^Sr (0.70806-0.70915) (Table S4). The highest value belongs to the LS419 *Equus sp*. The Sr isotope ratios for human teeth enamel and bones vary from 0.70822 to 0.70955, and from 0.70810 to 0.70894, respectively (Table S4). It is evident the difference between the isotope compositions of water samples and those of the biological material. In order to determine the bone preservation status and to verify the occurrence of post-depositional alteration FTIR analyses have been performed on 14 bones and, as an exploratory sample, one tooth (more details in Supplementary, chapter 3). LS151, LS418, and LS2176 have ^87^Sr/^86^Sr similar to the cave soil, while LS2177, LS2203, LS865, LS120, LS2208 have higher Sr isotope ratios (Tab S2 and Tab S3). Moreover, some of the analysed individuals have enamel teeth and bones with similar ^87^Sr/^86S^r (LS896, LS2203, LS2208, LS2176, and LS2177), while others - LS150, LS1956 and LS882 - have teeth and bone with different Sr isotope compositions (Fig. S4). Interestingly, all of them return similar FT-IR spectra (Fig. S5), showing an enlarged hump in the OH-stretching region overlapped by signals at ca. 2960 and 2880 cm^−1^ from ν(CH-)Lipids, a strong peak at ca. 1990-2000 cm^−1^ from cyanamide, and several absorption bands between ca. 1794 cm^−1^ and 400 cm^−1^. These characteristics can be assigned to the most important functional groups in bone, e.g., HPO_4_^2-^, PO_4_^3-^, CO_3_^2-^ and amide ^39–45^. Spectra are also characterized for an evident ν1(PO_4_^3-^) absorption peak at ca. 1045 cm^-1^ and the occurrence of fluorapatite (ca. 1090 cm^-1^), suggesting an expected (but not complete) deproteinization with reference to fresh (modern) bones ^43^. Since the young age ^46^, the LS 2176 infant lacks of a clear peak at 1415 cm^-1^ coupled to the 1450 cm^-1^ one (although a value can be assigned; see section 3 of Supplement). The C/P index, suitable both for pellets and DRIFT FT-IR techniques ^43^ is above 0.34 and significantly higher than 0.15, which is the considered threshold for altered bones. In addition, the order degree in phosphate minerals (CPI) varies between 2.7 and 3.4. Although these values could be slightly overestimated by DRIFT, they are generally lower than 3.4, which usually sets the limit between modern and archaeological bones. Therefore, the bones could not be considered diagenetically altered. Even though there are de-proteinized and de-phosphatized samples such as LS2208, LS151, LS120, LS2161, (Supplementary, Fig. S6), any contribution of type-A carbonate can be hypothesized except for the LS2176 bone^39^. Overall, the FT-IR indexes are not correlated with stable isotope compositions, whereas correlations between the C/P index and ^87^Sr/^86^Sr ratio (Supplementary, Fig S6d) are pointed out. These results, suggest that the ^87^Sr/^86^Sr teeth-bone pairs as reliable markers for hypothesizing the mobility pattern of people from La Sassa.

### Carbon and Nitrogen isotopic results

Forty human samples (87%) fitted the quality criteria related to collagen preservation (Table 1).

**Table 1:** Carbon and Nitrogen isotopic results for human bones.

| Samples | US | Bone | Chronology | $\delta^{13}\text{C}$ (‰) vs. VPDB | $\delta^{15}\text{N}$ (‰) vs. Air | %N | %C | C/N |
| --- | --- | --- | --- | --- | --- | --- | --- | --- |
| LS8 | Anfratto N | Femur L | CA | -19.9 | 8.9 | 13.1 | 40.4 | 3.6 |
| LS120 | 19 | Mandible | CA | -19.1 | 9.9 | 15.0 | 43.9 | 3.4 |
| LS121 | 19 | Radius R | CA | -20.0 | 8.3 | 13.7 | 39.3 | 3.3 |
| LS122 | 19 | Femur R | CA | -20.8 | 7.9 | 16.8 | 39.4 | 3.4 |
| LS126 | 19 | Femur L | CA | -19.9 | 8.7 | 14.1 | 42.1 | 3.3 |
| LS130 | 19 | Femur L | CA | -19.9 | 8.7 | 14.1 | 42.1 | 3.3 |
| LS132 | 19 | Radius R | CA | -19.6 | 9.2 | 12.4 | 35.2 | 3.3 |
| LS133 | 19 | Tibia L | CA | -21.0 | 9.5 | 15.0 | 44.0 | 3.3 |
| LS134 | 19 | Tibia R | CA | -20.0 | 9.4 | 15.2 | 42.2 | 3.2 |
| LS145 | 19 | Radius R | CA | -19.3 | 9.1 | 12.5 | 35.9 | 3.3 |
| LS146 | 19 | Tibia R | CA | -20.3 | 8.9 | 14.4 | 44.8 | 3.3 |
| LS150 | 19 | Mandible | CA | -19.7 | 9.8 | 13.9 | 43.7 | 3.6 |
| LS164 | 19 | Radius R | CA | -19.1 | 9.1 | 13.2 | 38.8 | 3.2 |
| LS212 | 19 | Mandible | CA | -19.3 | 9.8 | 16.4 | 48.3 | 3.4 |
| LS1956 | 19 | Mandible | CA | -19.6 | 9.4 | 17.2 | 54.0 | 3.6 |
| LS68 | 45 | Femur L | CA | -19.9 | 9.9 | 13.4 | 46.1 | 3.4 |
| LS565 | 55 | Mandible | CA | -20.7 | 8.4 | 13.9 | 41.7 | 3.5 |
| LS578 | 55 | Mandible | CA | -19.8 | 8.5 | 17.6 | 53.9 | 3.6 |
| LS585 | 55 | Tibia L | CA | -20.3 | 10.2 | 17.7 | 50.1 | 3.3 |
| LS892 | 75 | Tibia R | EBA | -21.7 | 8.2 | 13.6 | 40.5 | 3.4 |
| LS842 | 78 | Humerus R | EBA | -19.7 | 8.7 | 26.7 | 78.5 | 3.4 |
| LS 865 | 78 | Mandible | EBA | -20.5 | 8.7 | 13.3 | 41.1 | 3.6 |
| LS876 | 78 | Humerus R | EBA | -20.5 | 8.7 | 16.0 | 47.6 | 3.5 |
| LS878 | 78 | Tibia R | EBA | -21.6 | 8.5 | 13.9 | 43.3 | 3.6 |
| LS879 | 78 | Tibia L | EBA | -21.2 | 8.3 | 15.2 | 49.5 | 3.6 |
| LS882 | 78 | Mandible | EBA | -20.1 | 8.7 | 16.1 | 46.7 | 3.4 |
| LS889 | 78 | Humerus R | EBA | -20.7 | 8.6 | 17.6 | 53.8 | 3.6 |
| LS916 | 78 | Mandible | EBA | -20.3 | 7.4 | 12.3 | 38.4 | 3.6 |
| LS1978 | 78 | Mandible | EBA | -20.4 | 7.4 | 13.8 | 42.8 | 3.6 |
| LS2126 | 78 | Humerus R | EBA | -20.4 | 8.3 | 19.0 | 54.5 | 3.3 |
| LS2139 | 78 | Humerus R | EBA | -20.4 | 8.5 | 15.4 | 45.0 | 3.4 |
| LS2161 | 78 | Mandible | EBA | -20.7 | 8.2 | 12.2 | 37.1 | 3.5 |
| LS2168 | 78 | Humerus L | EBA | -20.5 | 7.9 | 17.0 | 49.1 | 3.4 |
| LS2177 | 78 | Mandible | EBA | -20.7 | 8.3 | 14.8 | 43.3 | 3.4 |
| LS2178 | 78 | Mandible | EBA | -20.7 | 8.3 | 12.6 | 38.0 | 3.5 |
| LS2203 | 78 | Mandible | EBA | -20.5 | 9.1 | 14.3 | 44.9 | 3.6 |
| LS2208 | 78 | Mandible | EBA | -20.5 | 9.1 | 13.1 | 41.3 | 3.5 |
| LS2403 | 97 | Mandible | CA | -20.5 | 8.6 | 15.3 | 47.6 | 3.6 |
| LS 4852 | 97 | Mandible | CA | -20.2 | 9.1 | 15.4 | 45.1 | 3.4 |
| LS2176 | RA | Femur L | MBA2 | -14.3 | 10.6 | 12.3 | 35.4 | 3.3 |
| LS151 | 19 | Femur R | CA | - | - | - | - | - |
| LS869 | 78 | Tibia R | EBA | - | - | - | - | - |
| LS907 | 75 | Tibia L | EBA | - | - | - | - | - |
| LS1959 | 55 | Tibia R | CA | - | - | - | - | - |
| LS1960 | 55 | Tibia R | CA | - | - | - | - | - |
| LS1961 | 55 | Tibia R | CA | - | - | - | - | - |

The isotope values for faunal remains have been analyzed to characterize the resources available as putative prey for people buried in La Sassa, despite most of the faunal remains should be dated to the BA (Table 2). The faunal δ^13^C and δ^15^N range between -22.5‰ to 20.1‰ and 4.7‰ to 9‰, respectively. Overall, these herbivore isotopic data are consistent with a terrestrial C3-based European ecosystem ^47^.

**Table 2:** Carbon and Nitrogen isotopic results for faunal bones.

| | US | Species | $\delta^{13}\text{C}$ (‰) vs. VPDB | $\delta^{15}\text{N}$ (‰) vs. Air | C/N |
| --- | --- | --- | --- | --- | --- |
| LS481 | 59 | *Bos taurus* | -20.5 | 9 | 3.3 |
| LS514 | 59 | *Sus domesticus* | -20.9 | 5.2 | 3.4 |
| LS793 | 49 | *Bos taurus* | -22.5 | 5.5 | 3.3 |
| LS595 | 55 | *Ovis vel capra* | -21.7 | 4.7 | 3.2 |
| LS535 | 49 | *Cervus elaphus* | -22.4 | 5.3 | 3.3 |
| LS794 | 49 | *Bos taurus* | -21.8 | 5.5 | 3.2 |
| LS517 | 59 | *Sus domesticus* | -21.2 | 4.9 | 3.3 |

Taking all 40 human individuals, δ^13^C varies from -14.3 to -21.7‰, while the δ^15^N values range between 7.4 to 10.6‰. Moreover, an alleged contribution from freshwater resources cannot be ruled out for most of the sample, while there is no clear indication of marine species consumption. The isotopic values of LS2176 (δ^15^N =14.3‰ and δ^13^C=10.6‰) clearly differ from the others (Fig. 3).

**Figure 3:**
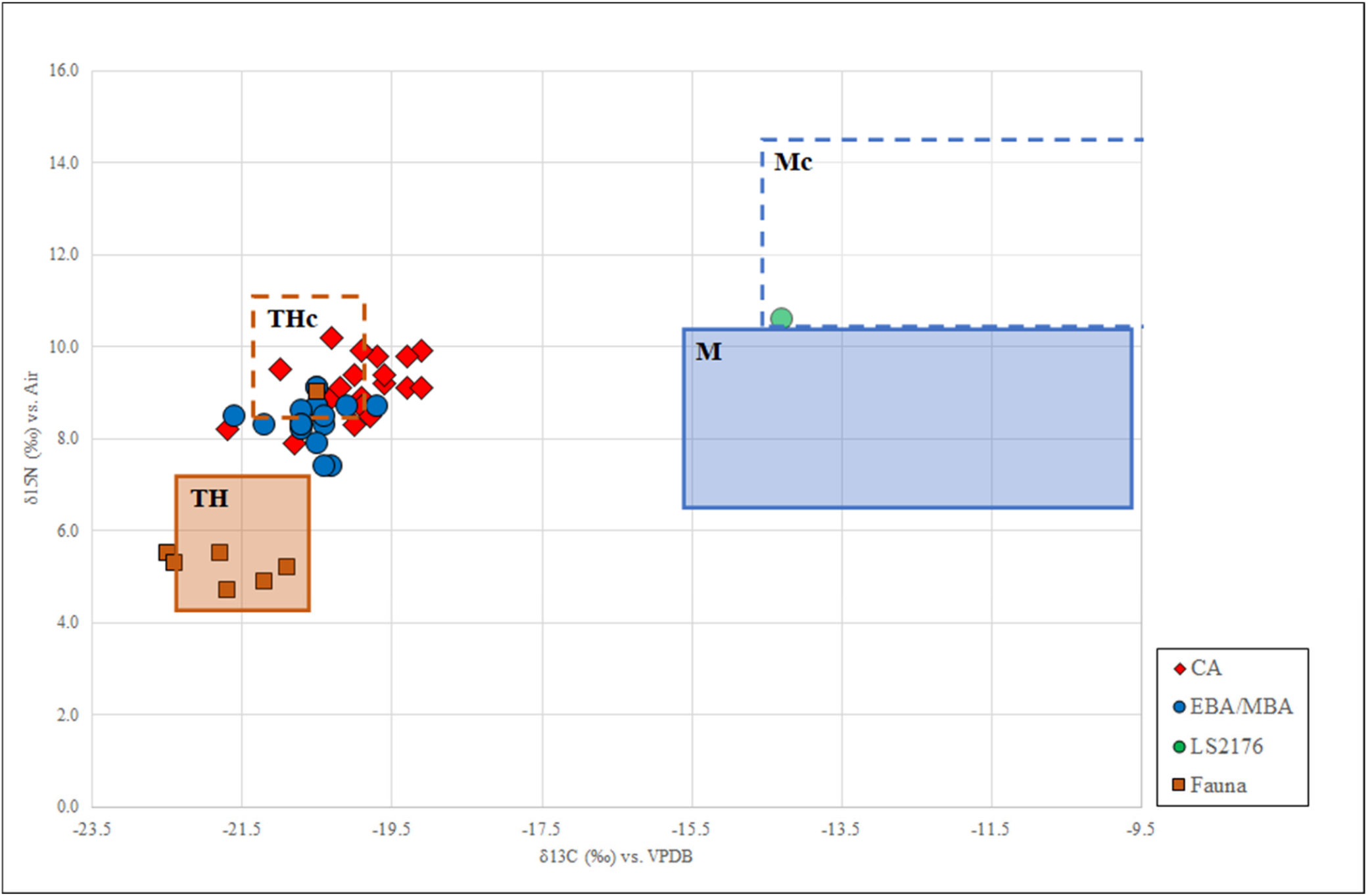
Plot of δ^13^C and δ^15^N values. The faunal data have been clustered according to their habitat and position in the trophic chain (rectangular boxes). The rectangular dotted line boxes are estimated from the faunal rectangular boxes, shifted by 1‰ on the δ^13^C axis and 4‰ on the δ^15^N axis to account for the trophic shift in isotope values from food source to consumer. TH and M: terrestrial herbivores and marine fauna, respectively. THc and Mc: terrestrial herbivore consumers, and marine resource consumers, respectively.

The faunal average values (δ^13^C= -21.5 ± 0.9 s.d. and δ^15^N = 5.7 ± 1.5 s.d.) were used to predict the local baseline of the trophic food chain. In figure 3 the data are compared to the human dietary reconstruction through the linear model proposed by Fraser et al. ^48^ and Fontanals-Coll et al. ^49^. All the analysed individuals roughly overlap the box of the terrestrial herbivore consumers (THc). The individual LS2176 is at odd, suggesting a different dietary model.

## Discussion

The topographical location of La Sassa and the chronology of its exploitation make it an outstanding context to explore the complex demographic scenario characterizing the CA/BA boundary in Central Italy. Despite the self-explanatory difficulties in dealing with chaotic bone assemblages, we leveraged multiple absolute radiocarbon dates to trace the different periods of cave exploitation. Accordingly, people recovered in Rooms 1 and 2 were dated to a roughly continuous phase starting in CA up to EBA. However, a funeral setting apart from the main Rooms returned the human remains of a toddler (LS2176, in the RA area) that is unrelated to the bone assemblages. The individual is chronologically apart from the others and dated as one of the latest individuals buried in the cave.

In order to catch up the biocultural features related to the communities using La Sassa as a funerary shelter, we chose to dissect the dietary habits through carbon and nitrogen stable isotopes. The present survey comprises the isotopic analysis of 40 human specimens recovered in La Sassa, representative for at least the whole sample identified, even though multiple districts could be redundant. The values gathered for bones recovered in Rooms 1 and 2 are consistent with a diet mainly grounded on C3 plant-based ecosystem (Fig. 3). The nitrogen signatures are consistent with a moderate protein intake, resulting from the consumption of local fauna resources. However, the chronological assessments showcase that people buried in the CA were characterized by significantly different isotopic signatures than people from the BA layers (Mann-Whitney δ^13^C U=66; p=0.0005; δ^15^N U=65; p=0.0005), prompting a slight change in dietary supply due to a drop in protein consumption through time. This shift towards lowest δ^15^N and slightly more negative δ^13^C values in the BA individuals is consistent with a fully grounded farming-derived diet following the tuning of the farming practices.

The data obtained from the CA layers of La Sassa are compared with roughly coeval central Italian contexts ^31^ (Supplementary Table S8). Significant differences for δ^13^C could be scored between La Sassa and Tuscans and Marche samples from Podere Cucule, Grotta del Fontino, and Fontenoce Recanati - even though they are all characterized by C3 based diet regime ^31^, while there are similarities with Le Lellere and Buca di Spaccasasso. However, the shallow δ^13^C values for all the sites led to exclude a staple consumption of C4 plants.

So far, La Sassa is significantly different from Le Lellere and Grotta del Fontino to what regards the nitrogen signature. However, as some referenced sites returned only a few individuals, these results suggest a subsistence affinity of the CA community buried in La Sassa and the other Central Italian communities, with some exceptions due to local preferential exploitation. The ecological association between the southern Tuscany and the Pontine plain is worth considering. Both the areas were characterized by coastal lakes that could be suggestive for similar diet, even accounting for only sporadic marine food assumption ^31^. However, La Sassa is slightly different from Grotta del Fontino (n=24; Mann-Whitney δ^13^C U=158; p=0.0329; δ^15^N U=154; p=0.0263), suggesting a different economic strategy. In this respect, the trade magnification rising in the CA could account for the local differences. Indeed, a trans-Apennine trade was previously advocated for addressing the similarities between the eastern Marche region and the southern area of Tuscany ^31,50,51^. The only southern Italian CA context with comparable data is Grotta di Donna Marsilia in the Calabria region^32^ where the unique human finding found therein fits the data gained for La Sassa CA people. If only a few studies are available for CA communities, a deeper knowledge concerning the Italian BA has been acquired recently through isotopic studies highlighting socio-economic transitions and new dietary habits grounded on the introduction of novel resources as the C4 plants ^29,30,32,34,52^.

The samples from BA layers from La Sassa are compared with BA sites from the Italian peninsula. The BA individuals from La Sassa isotopically match Ballabio sample (Supplementary Table S9) (n=22; Mann-Whitney δ^13^C U=128; p=0.0557; δ^15^N U=134.5; p=0.0858), a Northern area close to Como, characterized by low carbohydrate consumption and medium protein intake comprising primarily terrestrial resources typical of a C3 plant-derived environment and sporadic exploitation of freshwater resources ^52^. The heterogeneity in food exploitation in BA seems to be related to local dietary preferences, leading to differences among communities, even in similar ecological contexts.

So far, no direct evidence of C4 plant introduction seems to be recognizable in the Pontine plain in the initial phases of the BA.

However, LS2176 suggests a different scenario in the late phase of the frequentation of La Sassa. LS2176 was an infant (ca. 1-2 years) recovered apart from the other bone assemblages in the South-Western fringe of the cave. The radiometric dating (3165 ± 40 BP) sets this individual between the Middle Bronze Age sub-phase 2 (MBA2) and the Recent Bronze Age (RBA). However, as cremation was the exclusive ritual in the RBA, it seems to be traceable to MBA2, but no later than MBA sub-phase 3. The δ^13^C value reveals a surprising value (−14.3‰). According to the differential dating and spatial distribution, the toddler should not be related to the other BA people recovered in the cave. Unfortunately, the area where this infant was recovered cannot be further explored due to a massive landslide. Thus, to date, it represents the only individual dated to the MBA2 in La Sassa.

Still, LS2176 was probably be breastfed, therefore this odd δ^13^C value also reflects the values for the mother’s diet. Despite this finding, the frankly less negative carbon signature leads us to hypothesize that the community relied on different dietary habits. Even accounting for the well-known differential breastfeeding/weaning derived signature ^53,54^ where δ^15^N and δ^13^C exhibit decreases exponentially with age and the loss of one trophic level (ca. 4‰ and ca. 1‰, respectively), we could associate the LS2176 shift to dietary habit experienced by the mother. Even though we cannot exclude the leveraging of, at least moderate, consumption of marine resources coming from the tidal environment, it is worth considering that LS2176 falls at the edge of the variability identified for consumers of marine resources. Additionally, the exclusive tidal resources consumption seems unreliable, even accounting for the nearest seashore, which is located at least 15 km away, with the Ausoni Mounts in between. Reasonably, the odd signature could derive from the exploitation of C4 plants. There are no available isotopic values for C4 plants in Italian prehistory to the best of our knowledge. Some historical millet and sorghum samples from African areas ^55^ represent the few available comparisons for exploring the C4 isotopic signatures, showing enriched δ^13^C values ranging from - 13.1‰ to -10.6‰. Accordingly, we cannot test whether the value recorded for LS2176 could be derived reliably from an exclusive consumption of these resources. However, it is worth mentioning that some Italian BA communities diets that are convincingly considered as grounded on a stable C4 plant dietary intake ^26,29^ pointed out carbon signatures comparable with those determined for LS2176 (δ^13^C = -14.9 ± 1.1‰ for δ^13^C at Olmo di Nogara, -15.2 ± 2.4‰ at Bovolone, and 15.4‰ at Dossetto di Nogara, -16.5‰ the less negative result at Misa cave, dated to the MBA).

So far, we cannot exclude that the isotopic signature for LS2176 could be derived from a mixed parental diet, as the tidal resources and C4 ones may not be mutually exclusive. However, whatever being the primary edible resources consumed by LS2176 and his/her nurse, it seems remarkable that the people visiting La Sassa cave in the late phases of its exploitation explored different dietary habits than the communities that established the funerary setting in the previous stages.

Certainly, the recovery of further coeval individuals could be meaningful in strengthening our interpretation. This finding represents the first documented individual clearly showing the occurrence of a C4-based diet in the Pontine area.

It seems to be remarkable that this novel diet occurred in people dated after the evidence from Misa cave (Latium)^29^, suggesting a spatial and chronological cline for the dispersal of the C4-based diet from North-Eastern Italy^34^ to the Southern peninsular areas. Moreover, to date, there has been a lack of evidence for the spread of C4 plant consumption in Southern areas, even though no late MBA sites were isotopically investigated ^32^.

The southward dispersal of human groups sharing a C4-based diet is in agreement with the genomic and cultural evidence. Indeed, human groups with a different genetic makeup seem to get through the Pontine plain and the surrounding areas in the MBA, as demonstrated by the different genomic characteristics between CA people buried in La Sassa and BA individuals from the neighbouring Grotta Regina Margherita^4^. Similarly, stylistic influences from the North are recorded in the La Sassa ceramic assemblage for the first time.

The dissection of the mobility pattern contributes to characterize people from La Sassa, showing that they could have spent their lives around the cave. The ^87^Sr/^86^Sr ratios between CA and EBA samples do not allow for identifying significant differences in teeth enamel, even though the variation rises in EBA. Conversely, the Sr ratios for bones is significantly different (median CA_teeth_: 0.7091, IQR CA _teeth_: 0.0004; median EBA _teeth_: 0.7089, IQR EBA _teeth_: 0.0006; median CA_bones_: 0.7083; IQR CA_bones_: 0.0002; median EBA_bones_: 0.7085; IQR EBA _bones_: 0.0003; Mann-Whitney U=49.5; p=0.24 for teeth enamel and U=25 p=0.042 for bones), suggesting a slightly dissimilar mobility pattern between the differential periods.

Overall, EBA data are more scattered than CA values, supporting the few isotopic studies about mobility for Italian prehistoric communities, suggestive for the development of a mobile behaviour starting from the BA ^25–27^.

Remarkably, human data seem to be unrelated to the hydrographic net identified by the sampled springs, suggesting that people exploited water sources other than those nowadays flowing around La Sassa, or that unsampled springs were exploited. These springs could be possibly fed by water isotopically different from the exploitation area. Indeed, thermal waters in the aquifers of Mt. Massico ridge, at the south of Mt. Aurunci, are featured by variable Sr ratio, ranging from 0.7080 to 0.7096) ^56^, because the isotopic signatures of the water are impacted also by the rocks of the catchment area, which could be isotopically different from those in the exploitation area ^57^. Accordingly, the signature for humans was compared to the isotopic fingerprint of multiple soils. The isotope variability of soils prevents for identifying some specific areas where the geographical origin of both CA and EBA communities could be hypothesized. The overall drop in ^87^Sr/^86^Sr between teeth and bones suggests a certain degree of mobility for people between childhood and adulthood (Fig. 4). Therefore, people buried in La Sassa spent their childhood elsewhere from the area surrounding the cave. Whatever being the difference between teeth and bone values, it should be noted that the samples do not return isotopic fingerprint matching that obtained for the cave soil (Supplementary Fig. S4). This evidence suggests that the dwellings for both CA and EBA people were established in areas other than La Sassa geological setting. Accordingly, we can assume that the sampled individuals lived away from La Sassa cave, and they did not depend upon the sampled hydrographic net. However, some individuals return similar signatures in teeth and bones, suggesting a sedentary lifestyle rather than extensive mobility behaviours, scattering their values around 0.7085 and 0.7089. We cannot determine the exact geographic origin for CA and BA individuals analysed, as their isotopic values match those of wide areas around the cave, as suggested by the map of the distribution of the soils (Fig. 4). Overall, it seems reasonable that people moved around La Sassa in adulthood from sites characterized by soil, food, water etc. with slightly different isotopic ratios. This speculation can be corroborated considering the variability in the Sr isotope composition of a ca. 78 km^2^ area around the cave (corresponding to ca. 5 km-radius area from the cave, selected to characterize the local baseline), where the significant soils were geochemically characterized (Fig. 5 and Supplementary Table S3).

**Figure 4:**
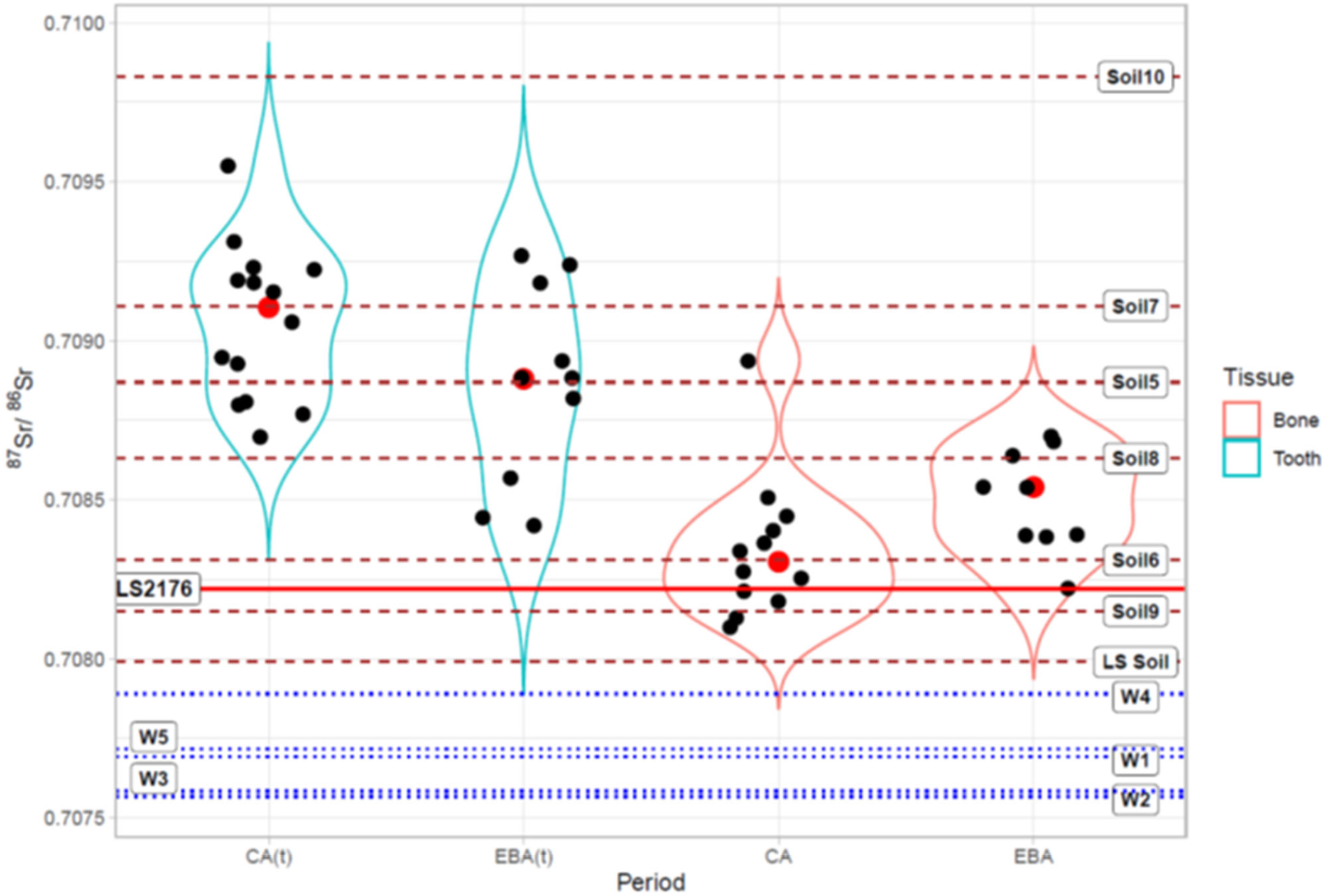
Violin plot for the Sr ratios (green for teeth (t), brown for bones) for CA and EBA individuals. LS2176 is represent by a continuous red line. Black dots represent single samples, red dots are the medians. The dashed lines represent the soils, while the dotted lines indicate the water samples.

**Figure 5:**
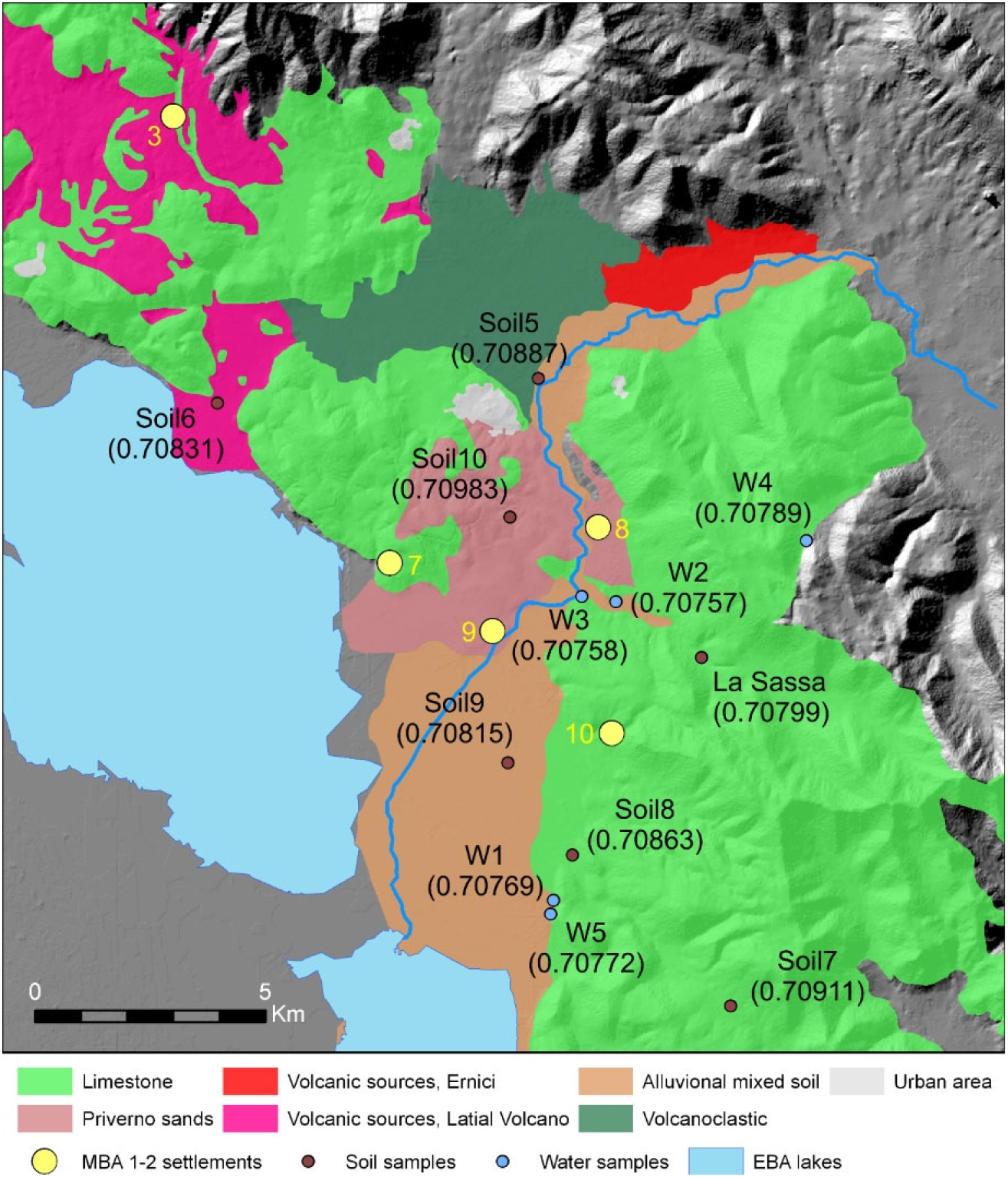
major rock and soil types around La Sassa cave. For the settlement numbers see figure 2. Background DEM from TINITALY/01 ^20^. W stands for Water sample.

However, some limitations should be considered in the dissection of the mobility pattern. Indeed, the Sr isotope values obtained for both CA and EBA individuals may be consistent with multiple communities sharing the environment around the cave, and we do not know if they were synchronic human groups. Even considering the single period – CA or EBA –, the value variation around the median suggests that only some individuals could have shared the same geochemical environment. These findings lead us to not rule out the hypothesis that multiple communities living in different places could have picked La Sassa as a funerary shelter through time. Indeed, it is also plausible considering the radiometric evidence concerning the cave’s prolonged exploitation, which could be consistent with a discrete selection of La Sassa as a funerary or ritual environment by people scattered throughout the area.

Unfortunately, no CA and EBA settlements are already discovered in the area, failing to support the geographic origin and demographic background of people buried in the cave. LS2176 ^87^Sr/^86^Sr fingerprint relies upon the local isotopic baseline. The paucity of the MBA anthropological finding in La Sassa prevents further speculations, even though its ratio seems to be in agreement with the data obtained for soils around already known MBA settlements around one hour walk from the cave (Supplementary, Fig. S10).

Even though we cannot ascertain the selection criteria for burying people in La Sassa, it should be noted that the demographic selection would not be the leading one, as adults and children of both sexes were recovered in the cave. Remarkably, the genomic evidence pointed out a certain relatedness in a sub-sample of the individuals (Saupe et al., 2021), suggesting that some factors (e.g., familial legacy) could have driven the selection of people to be buried in La Sassa, accounting for the ^87^Sr/^86^Sr clustering of some individuals.

## Conclusions

La Sassa represented a funerary and cultic shelter for a long time, starting from the CA up to the MBA. The recovery of human bones, faunal remains, and artifacts allows for evaluating the population dynamics occurring in the Pontine plain surrounding area. The isotopic evaluation shows that different communities could have exploited the cave in the CA and EBA. Unfortunately, the current archaeological landscape does not permit identifying the settlement areas for those human groups that experienced heterogeneous mobility behaviours and pick La Sassa as the funerary area for selected individuals.

CA and EBA individuals based their subsistence strategies upon a slightly different diet regime, even though both are grounded on terrestrial C3-based resources. However, La Sassa claims the arrival of human groups carrying a substantial innovation in the bio-cultural setting of the Central Italian communities. Indeed, the isotopic analysis identifies that people using La Sassa in the MBA2 grounded their diet upon different staple food, mainly represented by C4 plants. This evidence is consistent with the recent genomic findings tracing the arrival of people carrying a Steppe-related ancestry in Central Italy in MBA. Thus, our data confirm that the Pontine plain area was the stage for complex population dynamics. Indeed, the MBA1/MBA2 burden relates to a substantially different subsistence strategy. To date, the introduction of the C4 plant in the diet of peninsular Italian communities was seldom described, and the southernmost evidence was reported from a northern Latium funerary area.

For the first time ever, La Sassa highlights the southward spread of the C4 plant consumption, possibly related to the demic diffusion of allochthonous ancestries in the Central/Southern areas of the Italian peninsula. The southward both demic and dietary changes seem to have left bio-cultural traces in the Pontine area, allowing to define timely-based the arrival of these human groups through the dietary habits, the genomic features, and the cultural traits. Indeed, the ongoing extensive genomic evaluations will corroborate the arrival of people putatively carrying novel resources and ideas and hence new lifestyles in southern Italy following the MBA.

## Material and methods

### The archaeological excavation

In 2015, La Sassa cave was mapped and the surface artifacts collected. From 2016 to 2019, four excavation campaigns took place led by the Groningen Institute of Archaeology (Groningen University, Netherlands), in collaboration with the University of Rome Tor Vergata (Rome, Italy). Excavation and study permits have been received yearly from the Soprintendenza Archeologia, Belle Arti e Paesaggio per le Province di Frosinone, Latina e Rieti (2016: n. 4888 Class 34 31.C7/328.1; 2017: n. 9559 Class 34.31.07/74; 2018: n. 0013261-P Class 34.31.07/7.11.1/2018; 2019: n. 0016086-P Class 34.31.07/7.11.1/2018). The cave area has been virtually subdivided in progressively numbered rooms. Several soundings have been excavated in different portions of the cave. When the sounding area equals the room, the former has been labelled the same (Sounding Room 3); otherwise, an alphabetic code has been assigned to each sounding (sounding L, WD, SP, BA, all in Room 1) (Fig. 1b). The human and faunal remains have been individually collected and a progressive number with a prefix (LS) was assigned to each of them (LS1, LS2 etc). They have been recorded in a local coordinate system and then reprojected into the WGS84, UTM zone 33N (EPSG: 32633). The relative chronology is based on typo-chronological parallels and stratigraphic considerations, the absolute chronology is based on radiocarbon dates ^3,4^.

### Biological remains selection

The analysed skeletal series consists of over 800 human finds mainly recovered in Room 1 and Room 2 in a chaotic bone assemblage, probably due to taphonomic constraints, preventing the identification of individual burials. However, a comprehensive classification of each informative remain was performed according to bone or tooth, side, and, whenever possible, osteological markers suggestive for sex and age at death estimations.

The age of death was addressed by the wear of the teeth occlusal surface, the level of formation of the dental roots, and the synostosis degree of the tubular bones ^58,59^.

Sex identification was performed on dimorphic skeletal districts, mainly focusing on hipbones and skull traits ^60,61^.

The mandible was the most recovered anatomical district that allows for identifying at least 17 individuals (Minimum number of individuals; MNI). However, the joint information about sex, age at death, osteological characterization and osteometric analysis of the whole osteological material pushed us to consider a reliable sample consisting of at least 27 individuals.

Fifteen mandibles, a maxillary bone, six femur and 25 teeth were selected for tracing the ^87^Sr/^86^Sr ratio across the individuals. The selection was driven by macroscopic preservation (Supplementary Table S4). Due to the impossibility in identifying whether some anatomically unrelated bones pertain to a single individual, we selected 46 bones representative for the at least the whole sample identified to explore the carbon and nitrogen stable isotopes ratios, even though multiple districts could be redundant. To account for the ecological background, 7 faunal remains were also characterized isotopically (Table 2). To set the isotopic baseline within and around La Sassa cave, 6 groundwater and 7 soils samples were analysed for their Sr ratio (Supplementary Table S3).

### Sample preparation and analytical conditions for Sr isotope ratio determination

Teeth, bone, soil and water samples were prepared in the Clean Chemistry Laboratory of Istituto Nazionale di Geofisica e Vulcanologia, Sezione di Napoli Osservatorio Vesuviano (INGV, OV). Whole teeth were washed 2-3 times with Milli Q® H_2_O:H_2_O_2_=3:1 in an ultrasonic bath in order to remove organic material. Following this phase, a small piece was cut from each tooth with a dentist milling machine and Milli Q® H_2_O as a lubricant and washed again in an ultrasonic bath with a mixture of Milli Q® H_2_O:H_2_O_2_=3:1. Pieces of jawbones were washed in an ultrasonic bath 2-3 times with Milli Q® H_2_O at first, and then several times with a mixture of Milli Q® H_2_O:H_2_O_2_=3:1, in order to remove organic material. Once cleaned, samples were dissolved by using ultrapure HNO_3_ 65% in closed Savillex® vials. Following dissolution, the acid solution was dried down on a hot plate. The obtained solid fractions were re-dissolved in ultrapure 2.5N HCl and centrifuged for 10 minutes at 5000 rpm. Solutions were then loaded on quartz columns for the chemical separation of Sr by standard Sr isotope geochemistry procedures ^62,63^.

About 5 grams of soil samples from a radius of 5 km from the cave were left spinning in 100 ml of 1M ammonium acetate for 12 hours. Solutions were then filtered by using 0.45 microns filters, dried down on a hot plate, and dissolved in ultrapure 6N HCl at first, and 2.5N HCl at last. From the obtained, centrifuged acid solution, a 0.5 ml aliquot has been loaded on quartz columns for the chromatographic separation of Sr. Aliquot of groundwater, originating from the Apennine chain, and streaming in vicinity of La Sassa Cave, have been analysed with the aim of characterizing the local aquifers. Samples were dried down and the residues re-dissolved in ultrapure 2.5 N HCl. Before chemical separation of Sr, solutions were centrifuged for 10 minutes at 5000 rpm.

The Sr fractions dissolved in diluted HNO_3_ have been loaded on previously degassed zone refined Rhenium filaments, to carry out the measurement of ^87^Sr/^86^Sr isotope ratios by thermal ionization mass spectrometry (TIMS) techniques at the Radiogenic Isotope Laboratory of the INGV, OV. Determinations were performed with a ThermoFinnigan Triton TI multicollector mass spectrometer running in static mode. Measured _87_Sr/_86_Sr ratios were normalized for within-run isotopic fractionation to ^86^Sr/^88^Sr=0.1194. For each single measurement, the average 2σmean, i.e., the standard error with N=180, was ±0.000009. The mean measured value of ^87^Sr/^86^Sr for the NIST-SRM 987 international standard was 0.71023±0.00002 (2σ, N=171); external reproducibility (2σ) during the period of measurements was calculated according to ^64^. The measured Sr isotope ratios were normalized to the recommended value of NIST-SRM 987 (^87^Sr/^86^Sr=0.71025).

### Carbon and nitrogen isotope analyses

The collagen extraction from bones was individually performed following Longin’s protocol modified by Brown and colleagues ^65^, which was also simultaneously applied to a modern bovine sample as a reference. In order to obtain an acceptable yield of collagen, the extraction was performed on about 500 mg of bone powder collected by drilling. A concentration step was also carried out for all the samples to enhance the collagen yield through 30 kDa Amicon ® Ultra-4 Centrifugal Filter Units with Ultracel® membranes. Each sample of collagen weighed 0.8-1.2 mg and was analyzed using an elemental analyzer isotope ratio mass spectrometer at the iCONa (isotope Carbon, Oxygen and Nitrogen Analysis) Laboratory of the University of Campania. Carbon (δ^13^C) and nitrogen (δ^15^N) stable isotope ratios were measured in a single run on a Delta V Advantage isotope ratio mass spectrometer coupled to a Flash 1112 Elemental Analyser via a Conflow III interface (Thermo Scientific Milan, Italy). Results were expressed in δ notation ^66^ and reported in permille units. The measurements of δ^13^C were calibrated to the international standard VPDB with the standard reference materials IAEA-CH3, IAEA-CH6 and stable isotope ratio facility for environmental research at the University of Utah (SIRFER) yeast; δ^15^N measurements were calibrated to the international standard AIR with the standard reference materials USGS-34, IAEA-N-2 and SIRFER yeast. Typical analytical precision, evaluated from a repeated measurement of an internal standard, was 0.1‰ for δ^13^C and 0.2‰ for δ^15^N. The reliability of the procedure and the exclusion of exogenous contamination were accounted for through a comparison against established criteria to ascertain the percentages of carbon and nitrogen, atomic C/N ratios, and collagen yields ^67–69^.

The carbon and nitrogen contents and C/N ratio are listed in Table 1. To assess the preservation state of the extracted collagen, we considered carbon and nitrogen contents between 15-51% and 5-18%, respectively ^67^, and C/N ratios within the range 2.9 to 3.6 ^68^. The extraction yield was not used as a criterion ^67^ as the ultrafiltration technique was applied, and only samples with a yield of 0% were ruled out. Descriptive statistics and comparison tests were performed by R v.3.6.1 ^70^.

The linear mixing model proposed by Fraser et al. ^48^ and recently developed by Fontanals-Coll et al. ^49^ was implemented to detect the humans’ role compared to the available ecological resources. A theoretical terrestrial endpoint at -20.5‰ ± 0.9 was estimated (−21.5‰ adjusted to +1‰ for fractionation processes due to prey-predator relationship) from the mean δ^13^C value of terrestrial fauna. In the same way, a theoretical marine endpoint of about -12.6‰ ± 3.0 ^25,71,72^ was estimated. The theoretical thresholds for δ^15^N accounting for animal protein consumption were calculated adjusting for 4‰ the average faunal δ^15^N value (5.7‰± 1.5). All the above procedures resulted in a range of 8.2-11.2‰ for terrestrial faunal protein consumption in humans. Similarly, a theoretical marine threshold of about 10.5-14.5‰ was established.

### Sample preparation and analytical conditions for FT-IR analyses

FT-IR spectroscopy applies in investigation of human remains in archaeological and palaeontological contexts ^39–45,73^. Phosphate and carbonate minerals, as well as the collagen, that constitute bones and teeth can differ for structure and abundance based on freshness, aging maturation and post-mortem processes (Kramer and Shear, 1928; Rey et al., 1991). The presence and relative amount of important functional groups in bones, e.g., HPO_4_^2-^, PO_4_^3-^, CO_3_^2-^, and amide, can be defined by the various FT-IR, ATR, DRIFT or pellets, techniques ^43,e.g. 74^. Carbonate type A, carbonate type B, inorganic calcite, proteins, water, phosphate and fluoroapatite, can be identified and their relative abundance established. Therefore, the FT-IR spectra provide a simple screening about the state of conservation of proteins and the in vivo and post-mortem crystallizations in bones and teeth, as well as about structural resemblances.

FT-IR was used to investigate biominerals and collagen preservation (Beasley et al., 2014). Carbonate type A, carbonate type B, inorganic calcite, proteins, water, phosphate, and fluoroapatite were were identified and their relative abundance established by means of the diffuse reflectance infrared spectroscopy (DRIFT-FT-IR) at the Istituto Nazionale di Geofisica e Vulcanologia, Sezione di Napoli Osservatorio Vesuviano (INGV, OV) (Naples, Italy). The instrument used was a Nicolet 670 NexusTM equipped with a DRIFT accessory both by ThermoFisher Scientific S.p.a. The configuration included a heated ceramic (Globar) source, 670 Laser unit, KBr beamsplitter and an MCT detector. Samples were mixed and powdered with KBr in an agate mortar in a ratio of 1:10. Spectra were recorded on the mixture following the background attainment on the KBr alone at 100 scans and a resolution of 8 in the 5000–400 cm^−1^ range. For data acquisition and interpretation, we used the OMNIC Data Collector 5.2© software. Some samples have been done in duplicate or triplicate to assess analysis reproducibility. The errors in absorbance are on the order of 0.003; the various amine bands in the 1600 – 1450 cm^-1^ region are weak and caused largest uncertainty in absorption evaluation. In applying DRIFT-FT-IR technique, we careful consider the literature considerations about possible uncertainties compared to ATR and pellets ^43^. The indices in Table 3 were calculated by using the line between 1900-900 and 800-500 (750-400) cm^-1 39,41,43^. The relative amount of carbonate vs. phosphate (CC/PP) was also calculated from the ratio between the sum of absorbances at 1460 and 1425 cm^-1^ and the sum of absorbances at 605 and 568 cm^-1^ by using the baseline defined between 500 and 2000 cm^-1 40^: (A_1460_+A_1425_)/(A_605_+A_568_). Errors on index have been determined considering differences among samples analyzed in duplicate or triplicate.

**Table 3:** Infrared indexes with their significance and formula.

|  |  |
| --- | --- |
| BPI | Relative abundance of type B carbonate to phosphate: absorbance at $1415\text{ cm}^{-1}$ divided by the absorbance at $605\text{ cm}^{-1}$ ($A_{1415}/A_{605}$) |
| API | Relative abundance of type A carbonate to phosphate: absorbance at $1540\text{ cm}^{-1}$ divided by the absorbance at $605\text{ cm}^{-1}$ ($A_{1540}/A_{605}$) |
| BAI | Relative amount of B to A carbonate: absorbance at 1415 cm<sup>-1</sup> divided by the absorbance at 1540 cm<sup>-1</sup> ($A_{1415}/A_{1540}$) |
| PCI | Crystallinity: sum of the absorbances of the phosphate bands at 565 cm<sup>-1</sup> and 605 cm<sup>-1</sup> divided by the absorbance at 590 cm<sup>-1</sup> ($(A_{565}+A_{605})/(A_{590})$); |
| C/P | Relative amount of carbonate vs. phosphate: absorbance at 1415 cm<sup>-1</sup> divided by absorbance at 1035 cm<sup>-1</sup> ($A_{1415}/A_{1035}$) |
| Am/P | Collagen content of bones: absorbance of organic component at 1640 cm<sup>-1</sup> divided by the absorbance of $\nu_1\text{PO}_4$ at 1035 cm<sup>-1</sup> vibrational mode ($A_{1640}/A_{1035}$) |

## Authors’ contributions

MR, conceptualization, investigation, writing original draft, review, visualization; IA, conceptualization, investigation, writing original draft, review, visualization; MDV, OR, and CL review; MRDC carbon and nitrogen isotopic analysis, MFR, resources, data curation, review; MP, conceptualization, investigation, writing original draft, review, visualization; JS, conceptualization, investigation, writing original draft, review; FDA, conceptualization, investigation, writing original draft, review; LA, conceptualization, investigation, writing original draft, visualization, review, supervision, project administration, funding acquisition. All authors were involved in the interpretation of the results and contributed to the final version of the manuscript. The authors acknowledge Daniel F. Levey for the revision of the language throughout the manuscript.

## Availability of data and material

All data needed to evaluate the conclusions in the paper are present in the paper and the Supplementary Materials. The finds from La Sassa cave are stored in the Archaeology Laboratory of the University of Tor Vergata, Department of History, Culture and Society.

